# Epidermis Derived Lactate Promotes Sterile Inflammation by Inducing Metabolic Rewiring in Macrophages

**DOI:** 10.1101/2022.07.29.501937

**Authors:** Uttkarsh Ayyangar, Aneesh Karkhanis, Heather Tay, Aliya Farissa Binte Afandi, Oindrila Bhattacharjee, Lee Sze Han, James Chan, Srikala Raghavan

## Abstract

Dysregulated macrophage responses and changes in tissue metabolism are hallmarks of chronic, sterile inflammation. However, the metabolic cues that direct and support macrophage functions are poorly understood. Here, we show that during sterile inflammation in skin, the epidermis and macrophages uniquely depend on glycolysis and TCA cycle, respectively. This compartment separation is initiated by HIF1a stabilization and enhanced glycolysis in the epidermis. The end product of glycolysis, lactate is exported and utilized by the dermal macrophages to drive their effector functions. Notably, inhibition of lactate mediated crosstalk between the epidermis and macrophages leads to inhibition of sterile inflammation. Overall, our study identifies an essential role for the metabolite lactate in regulating macrophage response that can be effectively targeted to treat skin disorders such as psoriasis.

**One-Sentence Summary:** Epidermis derived lactic acid drives sterile inflammation by augmenting pro-remodeling state in dermal macrophages.

## Main Text

Inflammation occurring in the absence of external pathogens and tissue barrier breach is termed as sterile inflammation(*1*, *2*). Inability of the tissues to resolve sterile inflammation leads to progression of diseases such as cancer and rheumatoid arthritis(*1*). Tissue resident and recruited immune cells are key drivers of this inflammatory response(*2*). We previously reported that under sterile inflammatory conditions, the skin resident macrophages acquire an enhanced proremodeling fate that leads to degradation of the extracellular matrix(*3*). This alternative fate acquisition by the macrophages leads to progression of the diseased condition through exacerbated ECM disruption(*4*).

Interestingly, metabolic remodeling has been shown to complement fate decision in immune cells(*5*, *6*). In this regard, recent studies suggest that macrophages exhibit high degree of metabolic plasticity to fulfil their effector functions(*7*–*9*). Conceivably, in tissues, the metabolic pathways acquired by the macrophages must depend on exchange of nutrient sources with the macrophage-niche cells(*10*). The role of the macrophage-niche derived metabolites in regulating macrophage intrinsic metabolism, and in turn, effector functions, is poorly understood. In this report, using epidermal integrin beta1 KO mice and imiquimod induced mice model of psoriasis, we uncover a metabolic principle that drives pro-remodeling fate switch in macrophages in sterile inflammation.

As reported previously, loss of integrin beta1 (itgβ1) from the epidermal compartment in mouse embryonic skin, leads to augmentation of an inflammatory response characterized by increased macrophage cell infiltration and pro-remodeling fate acquisition(*3*, *4*). To gain insights into the metabolic states of macrophages and its niche cells, we used a previously generated NGS data from the epidermis, macrophages and fibroblasts of WT and itgβ1cKO embryonic skin and identified differentially expressed metabolic pathways in cKO epidermis(*4*). Pathway analysis and qPCR validation suggested an increase in expression of genes associated with the glucose uptake and glycolysis metabolism in the cKO epidermis (fig. S1A-C). Immunostaining analysis of the cKO skin with GLUT1 (Glucose transporter 1) and LDHa (Lactate dehydrogenase A) and western blot analysis of HK2 (Hexokinase 2) further corroborated these results (Fig. 1A-D). Notably, temporal analysis of cKO epidermis suggested increase in membrane GLUT1 expression at E18.5 (fig S1,D).

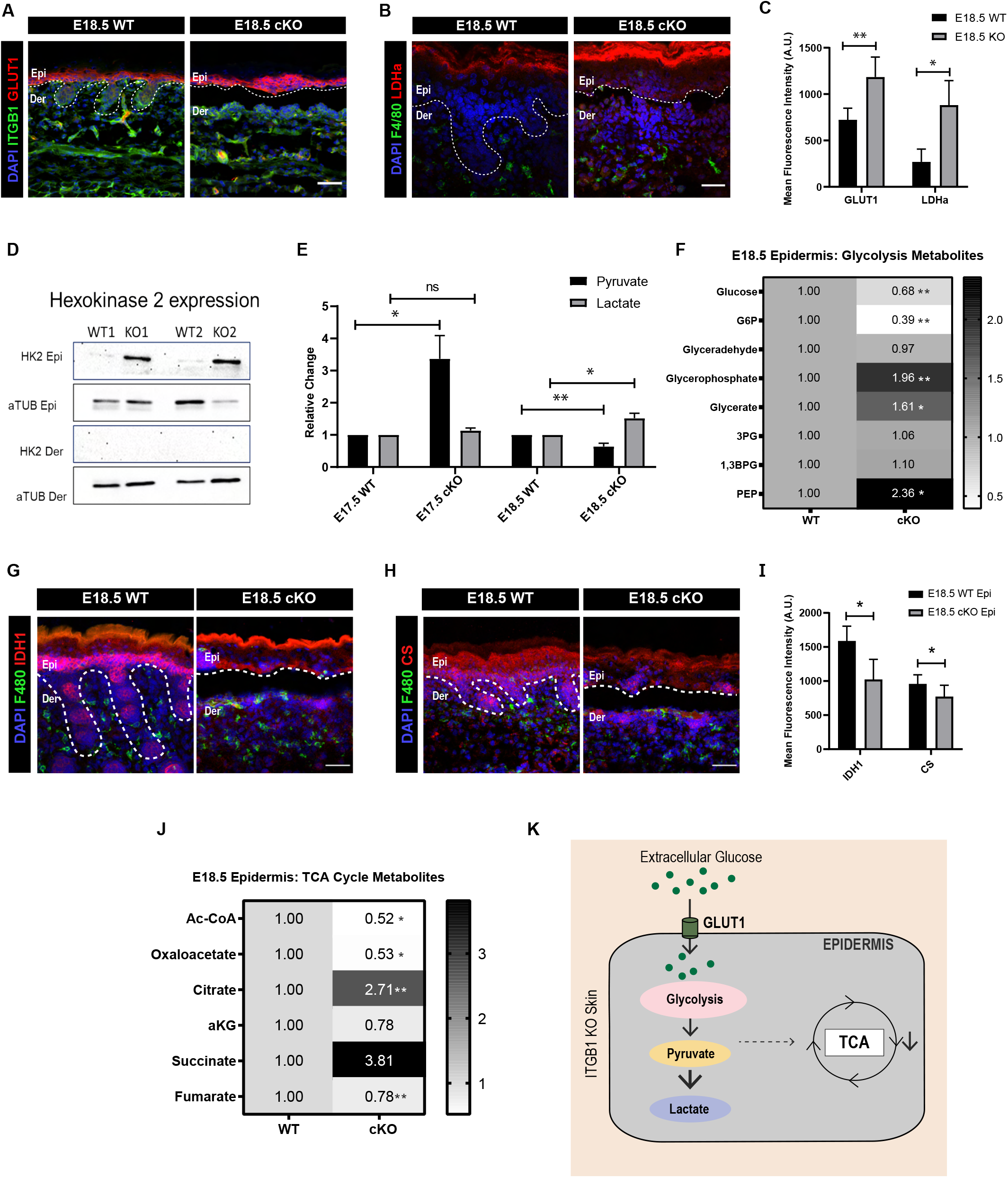

Glycolytic end products pyruvate and lactate feed directly into the tri-carboxylic acid cycle (TCA cycle) as part of central carbon respiration chain. We examined the expression of TCA cycle genes in the epidermal compartment from WT and cKO epidermis. Interestingly, NGS and qPCR analysis suggested a general reduction in all the TCA cycle enzyme transcripts in the cKO epidermis (fig. S1E,F). IDH1 (Isocitrate dehydrogenase 1) and CS (Citrate Synthase) immunostaining suggested a similar reduction in the TCA cycle enzyme expression cKO epidermis (Fig.1G-I). Furthermore, steady state metabolomics confirmed increase in glycolytic intermediate metabolites and glycolysis end product lactate, and decrease in TCA cycle metabolites in E18.5 cKO epidermis compared to WT (Fig 1E,F,J). Taken together, our results suggest augmentation of glycolysis that leads to enhanced lactate generation in the cKO the epidermal compartment (Fig 1K).

We next aimed to identify the molecular events that initiated the metabolic reprogramming in the cKO epidermal compartment. Hypoxia inducible factor (HIF1a) regulates glycolysis under both hypoxic and normoxic conditions (*11*). Notably, the NGS analysis suggested enrichment in pathways associated with response to hypoxia (fig. S2A). Consistent with this, we observed an increase in the expression of HIF1a and its downstream targets in the cKO epidermis compared to the WT (Fig. 2, A and B, fig. S2B,C,E). Temporal HIF1a expression analysis suggested augmentation of HIF1a expression in the KO skin as early as embryonic day E17.5 (Fig. 2A).

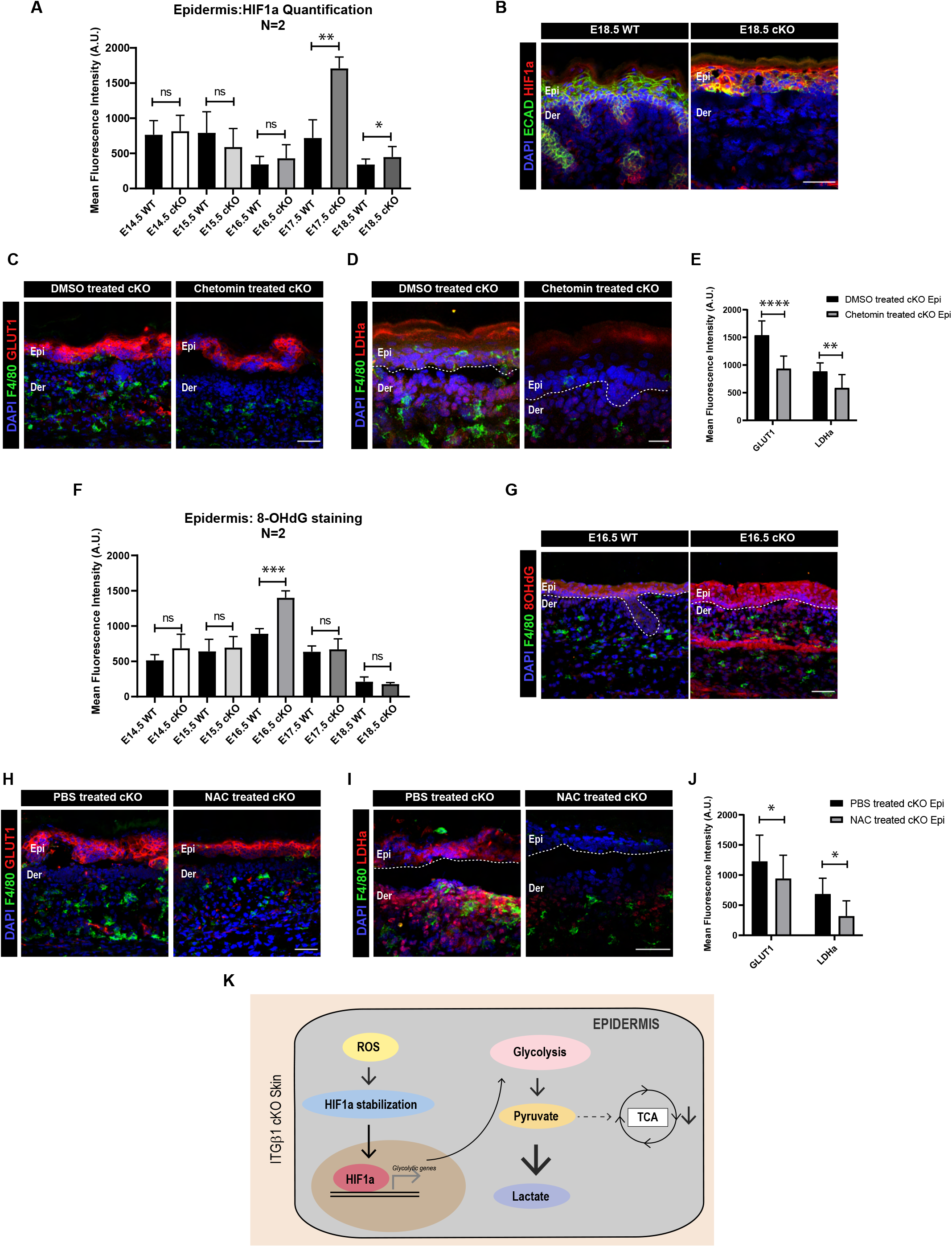

We next asked if increased HIF1a expression is directly associated with glycolytic upregulation in the epidermal compartment. To address this, we treated pregnant dams harbouring WT and cKO embryos with inhibitor of HIF1a dependent transcription, chetomin (3mg/kg)(*12*) (see Methods section, fig. S2G). The inhibition of HIF1a using chetomin not only reduced the expression of HIF1a targets – COX2 and KRT14, but also the expression of GLUT1 and LDHa in the in the cKO epidermis compared to DMSO treated cKO controls (Fig. 2C-E. fig. S2D,F). These results suggested that glycolytic programming in the epidermal compartment is augmented due to enhanced epidermal HIF1a expression and stabilization.

We next aimed to identify the mechanism of HIF1a stabilization in the cKO epidermis. Several studies have identified a role for reactive oxygen species (ROS) generated during trauma induced injury and tumors in stabilizing HIF1a response(*13*, *14*). Notably, GSEA analysis of the cKO epidermis suggested enrichment in pathways associated with an active response to increased ROS (fig. S3A). Temporal analysis of cKO and control skin using 8-OHdG (ROS indicator), suggested increase in ROS expression at embryonic day E16.5 cKO epidermis that quenches by E18.5 (Fig. 2F,G. fig. S3B). We reasoned that ROS burden in the KO skin might be counterbalanced by increased expression of ROS scavengers in the cKO epidermis. Consistent with this, we observed and validated an increase in the expression of ROS scavengers in E18.5 cKO epidemis (fig. S3C). This suggested that the upregulation of ROS in the cKO epidermis could be an early response to the loss of epidermal β1 which is eventually quenched through upregulation of ROS scavengers. We next attempted to identify possible sources of ROS in the epidermal compartment. qPCR analysis of different ROS sources suggested a global increase in genes associated with ROS generation through both oxidative and non-oxidative mechanisms (fig. S3C). This increase can be attributed to enhanced epidermal stress due to detachment from the extracellular matrix (ECM), the basement membrane. Notably, ROS increase upon ECM detachment has been reported previously in cancer cells during metastasis(*15*–*17*).

Since our temporal analysis suggested that ROS augmentation precedes HIF1a stabilization, we reasoned that ROS might be the potential activator for HIF1a signalling in the cKO epidermis. To address this, we treated pregnant dams harbouring WT and cKO embryos with the ROS scavenger, N-acetyl cysteine (NAC)(*18*) (fig. S2G). Interestingly, NAC treatment led to reduced expression of 8-OHdG and HIF1a in the cKO epidermal compartment compared to the PBS controls (fig. S3D,F). Immunostaining analysis further suggested a significant reduction in expression of HIF1a targets – KRT14, COX2, GLUT1 and LDHa in the cKO epidermis treated with NAC compared to the controls (Fig. 2H,I,J. fig S3E,G). The temporal analysis combined with pathway inhibition results suggest that an early ROS-HIF1a axis augments glycolytic metabolism in the cKO epidermis (Fig. 2K).

As described previously, enhanced glycolysis in the epidermal compartment led to lactate accumulation. We next aimed to understand the fate of epidermally generated lactate. Recent *in vitro* studies suggested that increase in lactate accumulation within the cells lead to increased membrane localization of lactate exporter MCT4 (monocarboxylic acid transporter 4)(*19*–*21*). Immunostaining of the cKO epidermis at E17.5 and E18.5 showed an increase in the membrane expression of MCT4 transporters (Fig. 3A). These data, in conjunction with metabolomics (Fig 1E), suggest enhanced lactate release from the cKO epidermal compartment.

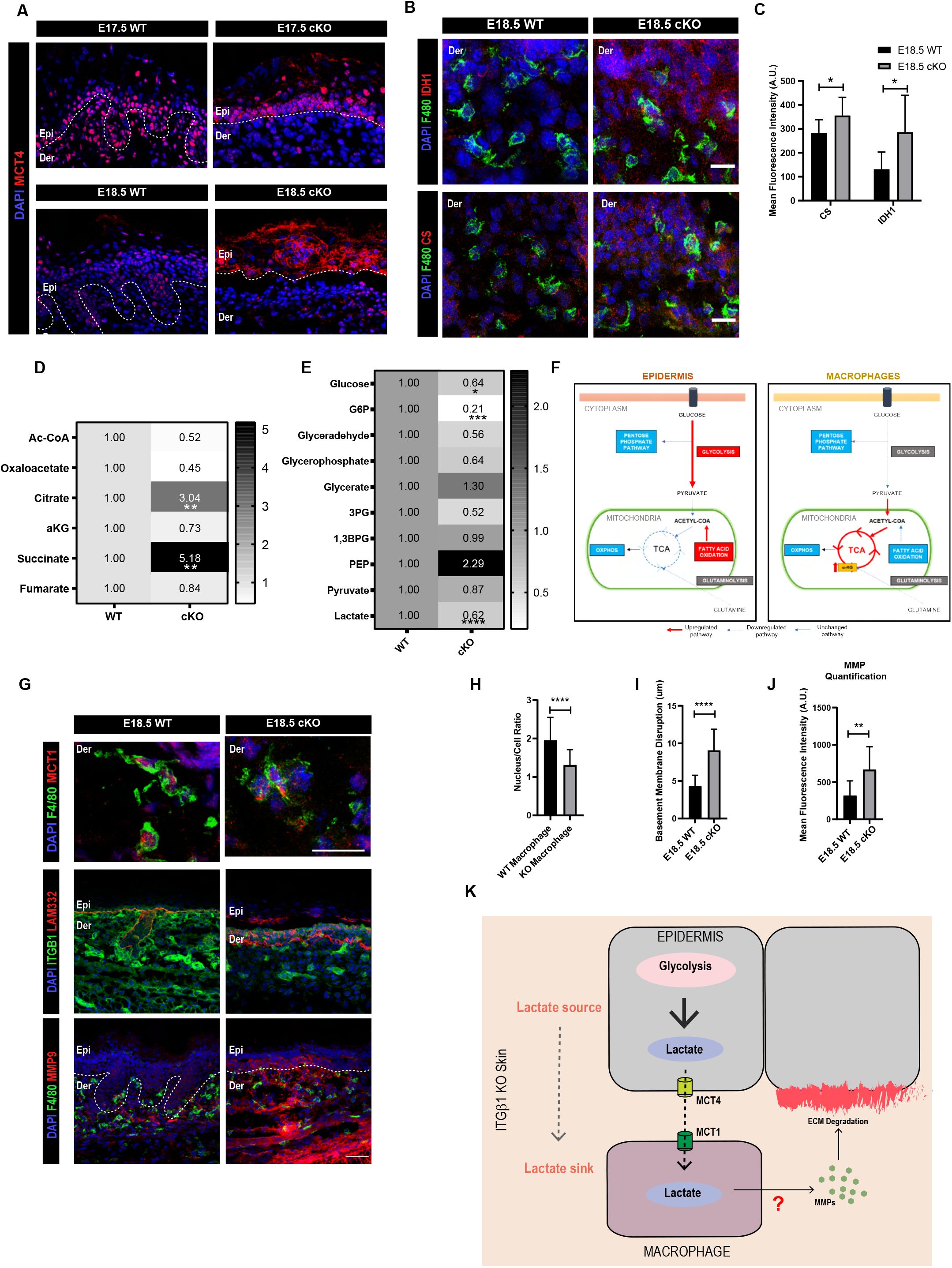

We therefore reasoned that the dermal fibroblasts and/or macrophages may serve as potential ‘sinks’ for epidermally derived lactate during sterile inflammation. NGS analysis of the macrophage compartment suggested an increase in the pathways associated with glucose deprivation (fig. S4A,B). This suggested a reduction in the glucose dependent metabolic program in the macrophages in the cKO skin. Notably, qPCR validation of the glycolytic genes from sorted macrophage population suggested no change in the genes associated with the glycolytic pathway (fig. S4C). The macrophages in the cKO skin did not express GLUT1 (fig. S4D). Interestingly, we observed and subsequently validated an increase in the genes associated with the TCA cycle in macrophages isolated from KO skin compared to the controls (fig. S4B,E). Immunostaining the F4/80+ve macrophages from the WT and itgβ1 cKO skin with TCA cycle enzymes – CS (citrate synthase) and IDH1 (Isocitrate Dehydrogenase 1) further corroborated the qPCR and NGS data (Fig. 3B,C).

Unlike the epidermal and macrophage compartment, we did not observe any change in the expression of enzymes associated with TCA cycle in CD45-ve fibroblasts (fig. S5A). This suggested that macrophages in the dermal compartment might be the key drivers of the TCA cycle. Notably, metabolomic analysis of the dermal compartment suggested a decrease in glycolysis metabolites and increase in TCA cycle metabolites. (Fig. 3D,E). Taken together, our analysis suggested that macrophages reduce dependence on glycolysis and instead augment TCA cycle in the KO skin. In addition, our results suggest a clear compartment-specific differentiation in glycolysis versus TCA cycle activity between the epidermis and macrophages, respectively, in the inflamed cKO skin (Fig. 3F).

We reasoned that since macrophages are decreasing dependence on the glycolytic metabolic pathway, they can utilize epidermally derived lactate as an alternative fuel to drive the TCA cycle (fig. S4F)(*22*, *23*). Immunostaining analysis suggested increased membrane localization of lactate importer, MCT1, in the macrophage compartment (Fig. 3G,H). This suggested that macrophages can act as potential sinks for lactate which in turn can be utilized to drive TCA in the macrophage compartment. Notably, membrane expression of MCT1 correlated with increased generation of MMP9 and basement membrane disruption (LAM332) in the KO skin underpinning a possible role for epidermally derived lactate in driving macrophage polarization during sterile inflammation (Fig. 3G,I,J).

Since both the epidermis and macrophage compartments in the itgβ1 cKO express lactate transporters, we reasoned that epidermally derived lactate might be sufficient to drive macrophage metabolic program and in turn its effector function. To test our hypothesis, we treated dams harbouring WT and cKO embryos with the MCT1 inhibitor, AZD3965(*24*) (fig. S2G). Interestingly, inhibition of lactate uptake by dermal macrophages in itgβ1 cKO animals led to a significant decrease in the expression of MMP9 which further resulted in a remarkable reduction in ECM degradation compared to DMSO treated controls (Fig. 4A-C). Additionally, inhibition of MCT4 (expressed in epidermis) using the MCT1/4 blocker, syrosingopine (SYRO)(*19*) with the identical treatment paradigm resulted in a similar reduction in the MMP9 generation and ECM degradation compared to the controls (Fig. 4A,D,E). These results underpin an important role for epidermal derived lactate in driving macrophage effector function during sterile inflammation in the KO skin. These results can further be extrapolated to sterile inflammatory disorders where increased lactate and MMP levels are associated with exacerbation of the diseased condition.

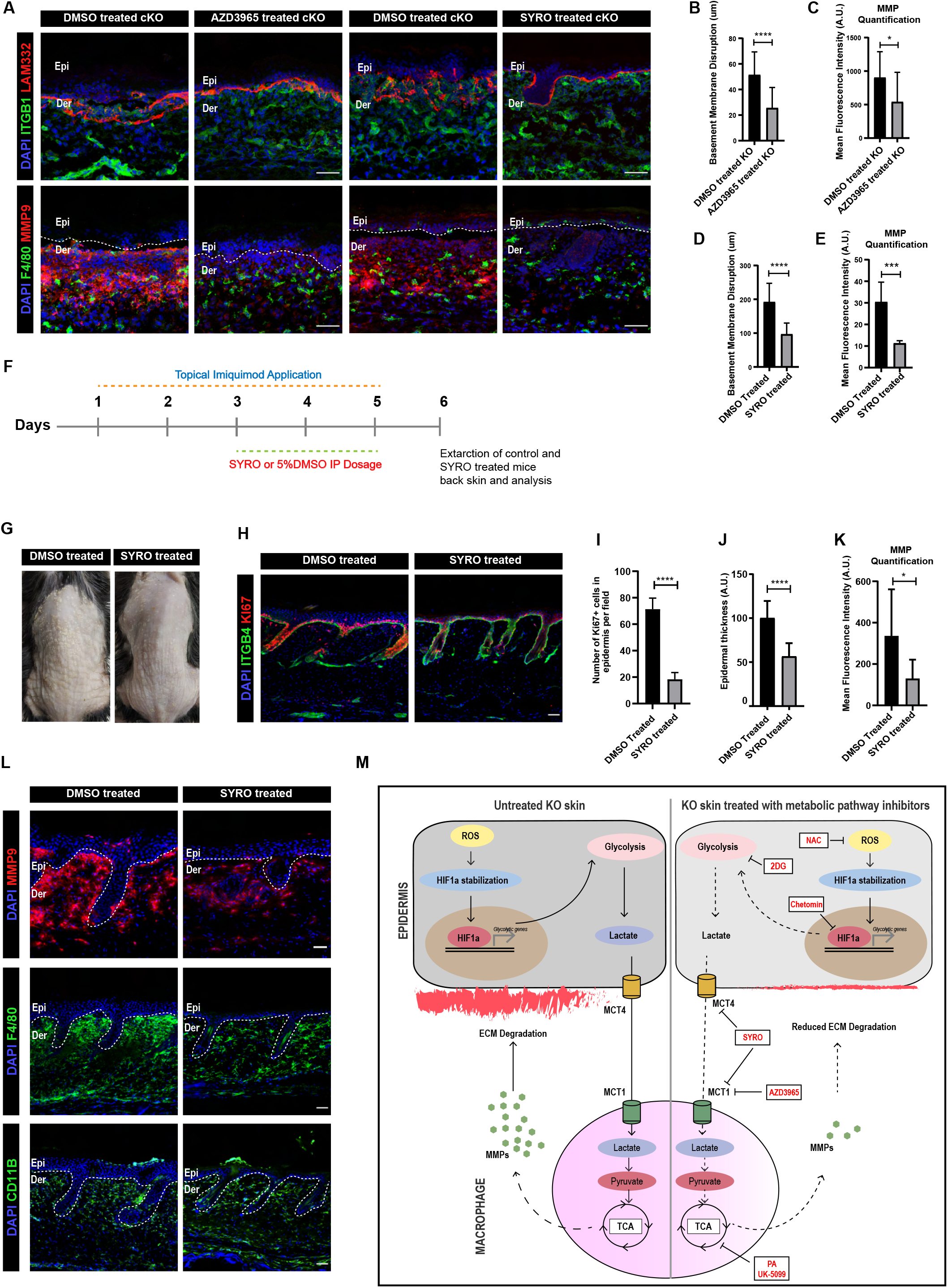

We next asked if the augmentation of the observed metabolic pathway in the macrophages was sufficient to cause basement membrane disruption in the KO skin. To address this, we treated pregnant dams with the small molecule inhibitors of the TCA cycle, pyromellitic acid (PA)(*25*), a fumarase inhibitor, and UK-5099(*26*), a pyruvate uptake inhibitor (fig. S2G). Interestingly, treatment with PA and UK5099 lead to a significant reduction in the basement membrane disruption and MMP9 expression in KO skin compared to the controls (fig. S6A-F). These results suggested that perturbation of macrophage intrinsic metabolic program was sufficient to inhibit macrophages effector function viz. MMP9 synthesis and concomitant basement membrane disruption itgβ1 cKO skin. We additionally established a positive correlation between lactate uptake and augmenting TCA cycle by macrophages in driving macrophage effector function.

To further strengthen our hypothesis, we treated pregnant dams with glycolysis inhibitor, 2DG (2 deoxy D glucose)(*27*) using identical treatment paradigm. We reasoned that since lactate transport inhibition led to reduction in macrophage pro-remodelling fate acquisition in the KO skin, inhibition of lactate source i.e., glycolysis in the epidermal compartment, should have similar consequences. Remarkably, blocking glycolysis with 2DG lead to a significant reduction in the generation of MMP9 and basement membrane disruption by macrophages (fig. S6A-C). On the other hand, inhibition of fatty acid metabolism, which is primarily augmented by the fibroblast compartment, using etomoxir(*28*) did not lead to reduction in macrophage mediated ECM degradation (fig. S5B,C). This suggested that the metabolic program augmented by the epidermal compartment, and not fibroblasts, have a direct consequence in regulating the effector functions of the dermal macrophages.

We asked if inhibition ROS and HIF1a, the upstream regulators or glycolysis, led to similar macrophage phenotype. We observed a significant reduction expression of MMP9 and extent of ECM disruption in NAC and chetomin treated KO skin compared to the controls (fig. S6G-K).

Finally, we aimed to understand the therapeutic applicability of our findings in the cKO skin. Previous studies on imiquimod induced mice model of psoriasis suggested increase in glucose utilization, glycolytic intermediates, and lactate in the epidermal compartment(*29*). We therefore hypothesized that lactate generated as a result of increased glycolysis in psoriatic epidermis might potentially drive psoriatic skin disease by augmenting fate changes in the macrophage compartment. Analysis of the mice skin post 5 days of imiquimod treatment suggested increased epidermal hyperproliferation and thickening which was concomitant with increased monocyte and macrophage burden (fig. S7A-D). In addition, we further observed increased MMP9 expression in the dermal compartment (fig.S7E-F). Notably, increased hyperproliferation in the epidermal compartment was associated with increased expression of GLUT1 and MCT4 (fig. S7G,I). This suggested that the epidermal compartment in the imiquimod induced mice model of psoriasis was a potential exporter of lactate. Additionally, macrophages show increased expression of TCA cycle enzymes, CS and IDH1 and lactate importer MCT1 (fig. S7H,J). This suggested that macrophages in the psoriatic skin potentially import lactate to drive TCA cycle which, in turn is necessary for their pro-remodelling fate switch. We finally asked if inhibition of lactate transport between epidermis and macrophages prevents macrophage polarization and in turn, psoriasis development in imiquimod induced mice model of psoriasis. Notably, treatment of mice with Syrosingopine lead to dramatic reduction in epidermal hyperproliferation, monocyte-macrophage burden and MMP9 expression in the psoriatic skin compared to the controls (Fig. 4F-K). These results establish lactate mediated epidermal – macrophage crosstalk as an important driver of the psoriatic skin disease.

Overall, using small molecule metabolic pathway inhibition studies, we show that perturbation of lactate transportation between epidermal and macrophage compartment and pathways leading to lactate synthesis leads to inhibition of macrophage pro-remodelling fate acquisition, thereby enhancing damage and inflammation in the KO skin (Fig. 4I).

In several skin disorders, the epidermal compartment has been shown to augment glycolytic program however, the functional consequence of this metabolic reprogramming is not well understood(*29*, *30*). Additionally, in sterile inflammatory diseases such as rheumatoid arthritis, while macrophages have been shown to be the associated with disease progression(*1*) the key molecular events that drive macrophages activation is not well understood. Our study suggests that under such conditions, glycolytic cell types may influence macrophage activation through enhanced lactate generation and release. Hence, lactate metabolism may be an important target to treat sterile inflammatory diseases.

Increased uptake of lactate by macrophages is important to drive their functional stares. Interestingly, recent studies have suggested that extra-cellular lactate can act as source of carbons for generating acetyl-CoA that changes acetylation states in tumour associated macrophages(*31*). In addition, TCA cycle metabolites can also be used as epigenetic adducts which can cause major changes in the transcriptional output of the immune cells(*32*). We speculate that lactate and TCA cycle intermediates might contribute to changes in the epigenetic landscape of macrophages that, in turn, may lead to enhanced MMP synthesis. The hypothesis needs further investigation.

Overall, our study identifies a lactate mediated crosstalk that drives sterile inflammation and psoriatic skin disease. The ability of lactate transport inhibitors to block the progression of psoriasis in our mouse models provides an exciting avenue to identify additional “druggable” metabolic pathways to treat sterile inflammatory diseases.

## Acknowledgments

We would like to thank Dr. Sunil Laxman (InStem, Bangalore), Dr. Ramanuj Dasgupta (Genome Institute of Singapore), and members of the Raghavan Lab for providing critical feedback on the work and manuscript. We thank the animal housing facility at NCBS/inStem.

## Author contributions

Conceptualization: UA, SR. Methodology: U.A., S.R., A.K., L.S.H. Investigation: U.A., H.T., A.F.B. Visualization: U.A. Funding acquisition: S.R. and U.A. Project administration: S.R. Supervision: S.R., L.S.H., J.C. Writing – original draft: U.A., S.R. Writing – review & editing: U.A., S.R.

## Data and materials availability

All data, code, and materials used in the analysis must be available in some form to any researcher for purposes of reproducing or extending the analysis. Include a note explaining any restrictions on materials, such as materials transfer agreements (MTAs). Note accession numbers to any data relating to the paper and deposited in a public database; include a brief description of the data set or model with the number. If all data are in the paper and supplementary materials, include the sentence “All data are available in the main text or the supplementary materials.”

