## Supplementary material for "Epidermis Derived Lactate Promotes Sterile Inflammation by Inducing Metabolic Rewiring in Macrophages": Materials and Methods

**Material and Methods**

**Animal Study**

Integrin β1 cKO animals (mixed background) were generated by crossing ITGβ1fl/+|KRT14-Cre males with ITGβ1fl/fl (C57B6J background) females. ITGβ1fl/+|KRT14-Cre males were generated by crossing KRT14-Cre homozygous males (CD1 background) with ITGβ1fl/fl (C57B6J background). Since integrin β1 cKO embryos are neonatally lethal, all the experiments reported in the current study have been done on embryos extracted from euthanized dams at specific embryonic stages.

Pregnant dams containing the KO and the WT embryos were housed at NCBS/inStem ACRC (Animal Care and Resource Centre) facility. Handling, breeding and euthanization of animals were done in accordance with the guidelines and procedures approved by the inStem IACUC (Institutional Animal Care and Use Committee). All experimental and breeder cages were maintained in SPF2 (Specific pathogen free 2) facility with standard ventilation, temperature (21 degree Celsius), 12-hour light and dark cycle, and sterilized food and water.

**Drug Treatments for β1 KO animals**

The pregnant dams containing the WT and integrin β1 cKO embryos were treated with small molecule inhibitors of specific metabolic pathways. All animals were treated for 3 days starting from E15.5. Embryos were extracted on E18.5 and analysed. In control experiments, pregnant dams were treated with the vehicles such as sterile PBS or 5% DMSO. The details of the drugs and treatment schedule is given in the table below:

| S.No. | Drug | Company | Catalogue Number | Target | Dosage | Dosage Route |
| --- | --- | --- | --- | --- | --- | --- |
| 1. | 2DG (2 Deoxy D Glucose) | Sigma | D8375 | Glycolysis | 1g/kg – concentration | Intraperitoneal |
| 2. | AZD3975 | MedChem | HY-12750 | MCT1 – Lactate Uptake Inhibitor | 200mg/kg – concentration | Intraperitoneal |
| 3. | Syrosingopine | Sigma | SML1908 | MCT1/MCT4 – Lactate transport inhibitor | 10mg/kg – concentration | Intraperitoneal |
| 4. | UK-5099 | Sigma | PZ0160-25MG | Mitochondrial Pyruvate carrier 1 inhibitor | 10mg/kg - concentration | Intraperitoneal |
| 5. | NAC | Sigma | A7250-10G | Reactive oxygen species inhibitor | 200mg/kg – concentration | Intraperitoneal |
| 6. | PA | Sigma | B4007 | Fumarase Inhibitor | 100mg/kg concentration | Intraperitoneal |
| 7. | Chetomin | Sigma | C9623 | HIF1a dependent transcription inhibitor | 3mg/kg - concentration | Intraperitoneal |
| 8. | Etomoxir | Sigma | E1905 | CPT1a (Fatty acid oxidation) inhibitor | 100mg/kg - concentration | Intraperitoneal |

**Imiquimod induced mice model for psoriasis**

C57B6/J mice back skin was shaved and 12.5 mg of commercially available 5% imiquimod (Glenmark) was applied topically daily on the shaved back skin. Vaseline was used as control for the above experiment. After 5 days of daily imiquimod or vasaline application, animals were euthanized as per guidelines and procedures approved by the inStem IACUC (Institutional Animal Care and Use Committee). The back skin was collected for further analysis. All experimental and breeder cages were maintained in SPF2 (Specific pathogen free 2) facility with standard ventilation, temperature (21 degree Celsius), 12-hour light and dark cycle, and sterilized food and water.

**Drug Treatments for Imiquimod animal models**

For lactate transport inhibition experiments, 12.5 mg of Imiquimod (Glenmark) was applied on the back skin of the mice induce psoriasis for a total of 5 days. From the third day onwards, animals were treated with intraperitoneal doses of Syrosingopine (SML-1908, SIGMA) at 10mg/kg concentration. The control animals were treated with 5% DMSO in sterile 1XPBS. Both male and female C57B6/J were used in these experiments. After 5 days, mice were euthanized as per guidelines and procedures approved by the inStem IACUC (Institutional Animal Care and Use Committee). The back skin was collected for further analysis. All experimental and breeder cages were maintained in SPF2 (Specific pathogen free 2) facility with standard ventilation, temperature (21 degree Celsius), 12-hour light and dark cycle, and sterilized food and water.

### Immunostaining

### Embryos extracted from euthanized pregnant dams were frozen in tissue freezing media (OCT) and 10-micron cryosections were collected on charged glass slides and stored in -80 ºC. For immunostaining, cryosections were thawed in room temperature (RT) for 5 minutes and fixed in acetone (Merck) for 5 minutes at -20 ºC or 4% paraformaldehyde (Sigma) at room temperature for 10 minutes. Paraformaldehyde fixed sections were permeabilized using permeabilization solution – 1XPBS plus 0.2-0.5% Triton X-100 (Sigma) for 10 minutes at RT. Fixed and permeabilized sections were blocked using 5%NDS (normal donkey serum, Abcam) in permeabilization solution. This was followed by addition of primary antibodies diluted in block. Details of the antibody dilutions and catalogue information is given in the table below. Primary antibody staining was carried overnight in 4ºC or 2 hours in RT. After washing with 1XPBS, secondary antibody staining was carried for 45 minutes at RT. Nucleus was stained using 1XDAPI (Sigma). The slides were covered with Moviol (Sigma), mounted and sealed. All images were taken in FV3000 5 Laser confocal microscope.

| S.No. | Antibody | Company | Catalogue Number | Dilution |
| --- | --- | --- | --- | --- |
| 1. | Integrin Beta1(ITGB1) | EMD Millipore | MAB1995 | 1:200 |
| 2. | Integrin Beta1(ITGB4) | BD Biosciences | [AB395027](http://antibodyregistry.org/AB_395027) | 1:200 |
| 3. | Glucose Transporter 1(GLUT1) | Abcam | AB115730 | 1:300 |
| 4. | Lactate Dehydrogenase alpha (LDHa) | Abcam | AB52488 | 1:200 |
| 5. | Hexokinase 2 (HK2) | Cell Signalling Technologiy | C64G5 | 1:1000 |
| 6. | F4/80 | Ebioscience | 14-4801-82 | 1:200 |
| 7. | Isocitrate Dehydrogenase 1 (IDH1) | Abcam | AB172964 | 1:300 |
| 8. | Citrate Synthase (CS) | Novus Biologicals | NBP2-13878 | 1:300 |
| 9. | Laminin 332 (LAM332) | Gift by Bob Burgeson | - | 1:500 |
| 10. | Matrix Metalloproteinase 9 (MMP9) | R & D Systems | AF909 | 1:50 |
| 11. | Hypoxia Inducible Factor 1 alpha (HIF1a) | Novus Biologicals | NB100-449 | 1:200 |
| 12. | 8 hydroxy guanine (8-OHdG) | Novus Biologicals | NB100-1508 | 1:200 |
| 13. | E cadherin (ECAD) | Thermo Fisher Scientific | 131900 | 1:500 |
| 14. | Keratin 14 (KRT14) | Abcam | AB181595 | 1:1000 |
| 15. | Cyclooxygenase 2 (COX2) | Abcam | AB15191 | 1:200 |
| 16. | Monocarboxylic acid transporter 4 (MCT4) | Sigma | HPA021451 | 1:200 |
| 17. | Monocarboxylic acid transporter 1 (MCT1) | Sigma | PA5-72957 | 1:100 |
| 18. | PECAM (CD31) | Ebioscience | 14031181 | 1:200 |
| 19. | Alpha-Tubulin | Sigma | T9026 | 1:1000 |

### Western Blotting

### Snap-frozen epidermis and dermis obtained from WT and KO skin were pulverized using sterilized pestles. Homogenized tissue was then suspended in RIPA lysis buffer containing 1Xprotease inhibitor cocktail. Protein extraction was facilitated using multiple freeze – thaw cycles followed by centrifugation at maximum speed for 15 minutes at 4^o^C. Protein concentration in the supernatant was measured using BCA assay (Promega). All protein isolate concentrations were normalised using RIPA-PIC buffer. 50ug of protein was loaded onto PAGE (8%) and electrophoresed, and transferred onto PVDF membrane (BioRad). Blocking of the membrane was done using 5%BSA (Sigma). Primary antibody staining was done overnight at 4^o^C. After washing with 0.1%TBST and secondary antibody (HRP conjugated) were added for an hour at RT. Unbound secondary antibodies were washed using 0.1%TBST and the blots were developed using ECL substrate (Thermo).

### RNA Extraction and Real Time PCR

### Total RNA was extracted using Trizol (Thermo) for epidermal tissue and Trizol LS (Thermo) for sorted fibroblast and macrophages. Visualization of RNA pellet was facilitated by using 1uL of Glycoblue (Thermo). Air-dried RNA was resuspended in nuclease free water (Thermo). Equal amounts of RNA obtained from KO and control skin compartments were used to prepare cDNA by using SSIII RT cDNA synthesis kit (Invitrogen). SYBR green (2X) master mix (Invitrogen) was used for real time PCR. Delta Ct method was used to quantify the relative changes in the level of transcripts. 18S was used as an endogenous control.

### The list of primers used is given below.

| Gene Name | Forward Primer | Reverse Primer |
| --- | --- | --- |
| Serpine1 | TTCAGCCCTTGCTTGCCTC | ACACTTTTACTCCGAAGTCGGT |
| Vegfa | GCACATAGAGAGAATGAGCTTCC | CTCCGCTCTGAACAAGGCT |
| Timp1 | GCAACTCGGACCTGGTCATAA | CGGCCCGTGATGAGAAACT |
| Fn1 | ATGTGGACCCCTCCTGATAGT | GCCCAGTGATTTCAGCAAAGG |
| Id2 | ATGAAAGCCTTCAGTCCGGTG | AGCAGACTCATCGGGTCGT |
| Krt14 | AGCGGCAAGAGTGAGATTTCT | CCTCCAGGTTATTCTCCAGGG |
| IGFBP3 | CCAGGAAACATCAGTGAGTCC | GGATGGAACTTGGAATCGGTCA |
| Gpx3 | TTTGTGCCTAATTTCCAGCTCTT | GTCCATCTTGACGTTGCTGAC |
| H6PD | ATGAAGCACACAGGCATTTGG | TCCAGGTATAGCTGAAACAGTCC |
| Nqo1 | AGGATGGGAGGTACTCGAATC | AGGCGTCCTTCCTTATATGCTA |
| Prdx6 | GTCGAGAAGGACGCTAACAAC | GGGTAGAGGATAGACAGCTTCAG |
| Alox5 | TTGCTCTCACAGTATGACTGGT | AGTATCCACGATCTGCTCGA |
| Alox8 | CTGTCAGCATCGTGGGAACC | GGAAGCGTCACCTCGAAGTC |
| Duox1 | AAAACACCAGGAACGGATTGT | AGAAGACATTGGGCTGTAGGG |
| DuoxA1 | ACCAAGCCAACCTTTCCAATG | GCCCCGATGAATAAGCTGGTC |
| Mpo | AGTTGTGCTGAGCTGTATGGA | CGGCTGCTTGAAGTAAAACAGG |
| Xdh | ATGACGAGGACAACGGTAGAT | TCATACTTGGAGATCATCACGGT |
| GLUT1 | CAGTTCGGCTATAACACTGGTG | GCCCCCGACAGAGAAGATG |
| Hk2 | TGATCGCCTGCTTATTCACGG | AACCGCCTAGAAATCTCCAGA |
| LDHa | CATTGTCAAGTACAGTCCACACT | TTCCAATTACTCGGTTTTTGGGA |
| PKM2 | GCCGCCTGGACATTGACTC | CCATGAGAGAAATTCAGCCGAG |
| MCT4 | TCACGGGTTTCTCCTACGC | GCCAAAGCGGTTCACACAC |
| Pfkfb3 | CCCAGAGCCGGGTACAGAA | GGGGAGTTGGTCAGCTTCG |
| ME2 | AAGGGAATGGCGTTTACGTTAC | GTACACAATCGGCATCAGACTT |
| Tpi | CCAGGAAGTTCTTCGTTGGGG | CAAAGTCGATGTAAGCGGTGG |
| Enolase 1 | TGCGTCCACTGGCATCTAC | CAGAGCAGGCGCAATAGTTTTA |
| Ogdh | GGAACTGCCCTCTAGGGAGA | GACGCTACCACTGTTAATGACC |
| FH1 | GAATGGCAAGCCAAAATTCCTT | CGTTCTGTAGCACCTCCAATCTT |
| Cs | GGACAATTTTCCAACCAATCTGC | TCGGTTCATTCCCTCTGCATA |
| Aco1 | AGAACCCATTTGCACACCTTG | AGCGTCCGTATCTTGAGTCCT |
| Aco2 | ATCGAGCGGGGAAAGACATAC | TGATGGTACAGCCACCTTAGG |
| IDH1 | ATGCAAGGAGATGAAATGACACG | GCATCACGATTCTCTATGCCTAA |
| IDH2 | CACCGTCCATCTCCACTACC | CAGCACTGACTGTCCCCAG |
| GOT2 | TGGGCGAGAACAATGAAGTGT | CCCAGGATGGTTTGGGCAG |
| SDHa | GGAACACTCCAAAAACAGACCT | CCACCACTGGGTATTGAGTAGAA |
| SDHb | GCTGCGTTCTTGCTGAGACA | ATCTCCTCCTTAGCTGTGGTT |
| SDHc | TGGTCAGACCCGCTTATGTG | GGTCCAGTGGAGAGATGCAG |
| SDHd | CGAAAGCGACATGGCGGTTC | GGTCCTGGAGAAATGCTGACAC |
| PFKl | GGAGGCGAGAACATCAAGCC | CGGCCTTCCCTCGTAGTGA |

### RNA Sequencing Analysis and data availability

### RNA sequencing used in the report has been done previously (Bhattacharjee et al., 2020). The data sets obtained from the report are submitted in NCBI with reference ID: SRP324814 (PRJNA739149). The data can additionally be accessed from the link: <https://dataview.ncbi.nlm.nih.gov/object/PRJNA739149?reviewer=1klobv7agr78somt2toldvkjvu>.

### Metabolomics

### 1. Material

### TCA cycle (ML0010) and glycolysis/gluconeogenesis (ML0013) metabolite libraries were obtained from Sigma-Aldrich (St. Louis, MO). Isotopic 13C15N-labelled amino acids (MSK-CAA-1) were obtained from Cambridge Isotope Laboratories (Andover, MA). Optima-LC/MS grade isopropanol (A461-4) and acetonitrile (A955-4) were obtained from ThermoFisher Scientific (Waltham, MA).

### 2. Methodology

### a. Tissue sample processing

### A total of 44 pre-weighed samples (22 epidermis, 22 dermis) were randomized and thawed prior to processing. To each sample, a volume of 300 µL of Type 1 ultrapure water was added and bead-homogenized at 4500 rpm, 3×10s cycles at 4°C on a Cryolys Evolution-Precellys Evolution tissue homogenizer (Rockville, MD). A volume of 100 µL of tissue homogenate was added to 500 µL ice-cold isopropanol containing 1.25 µM of 13C15N-labelled amino acids internal standards, and agitated for 1 hr to promote protein precipitation. Samples were then centrifuged at 20,000 g and the supernatant was collected. Extraction was repeated with another portion of 500 µL ice-cold isopropanol with internal standards. Pooled supernatants were then evaporated on a Labconco CentriVap concentrator (Kansas, MO). Samples were reconstituted in 100 µL acetonitrile:water (1:1), centrifuged at 14,000 g at 4°C for 10 min, then introduced into the liquid chromatography tandem mass spectrometry (LC-MS/MS) at injection volume of 1 µL.

### b. Liquid chromatography mass spectrometry

### Multiple reaction monitoring was performed on a Thermo Scientific Vanquish Duo UHPLC coupled to a Quantis TSQ Triple Quadrupole Mass Spectrometer. Chromatographic separation was done on a Waters Atlantis Premier BEH Z-HILIC column (1.7 µm, 2.1 mm × 100 mm) maintained at 45°C. Chromatographic separation was achieved at 0.45 mL/min using the gradient elution as follows: 0.0 min 99%B, 15.0 min 30%B, 17.9 min 30%B, 18.0 min 99%B, 25.0 min 99%B. Mass spectrometry was operated in electrospray ionization (ESI) mode with polarity switching, with source parameters as follows: spray voltage 3.5 kV (positive) -2.8 kV (negative), sheath gas 50 [arbitrary units], auxiliary gas 18 [arbitrary units], sweep gas 0 [arbitrary units], ion transfer tube temperature 325°C, vaporizer temperature 230°C. Details on mobile phases and compound-specific parameters are described in the Supplementary.

### Statistical Analysis

### All the statistical analysis presented in the paper were done in GraphPad Prism version 9.0.0. Two tailed students t test has been performed across all graphs with multiple biological replicates.
