## Supplementary material for "Epidermis Derived Lactate Promotes Sterile Inflammation by Inducing Metabolic Rewiring in Macrophages": Figure Legends

Fig. 1: Increased glycolysis and decreased TCA in epidermal compartment in β1 cKO skin

Epidermal compartment of cKO skin showing increased expression of GLUT1 (N=3) (A), LDHa (N=2) (B) quantified in (C). Western blot for Hexokinase 2 (hk2) in epidermal and dermal compartment of KO skin (D). Relative concentrations of pyruvate and lactate from E17.5 and E18.5 cKO epidermis (E). Glycolytic metabolites identified from epidermis of E18.5 β1 cKO and WT skin (F) (N=5). Reduced expression of CS (G), IDH1(H) and quantification (I) in epidermis of cKO compared to WT (N=2). Relative levels of TCA cycle metabolites in epidermal compartment of cKO and WT skin (J) (N=5). (K) Schematic showing glycolysis and TCA cycle changes as suggested by transcriptomic, proteomic and metabolomic approaches in the KO skin compared to WT. Scale bars: 50 µm. *p ≤ 0.05, **p ≤ 0.01, ***p ≤ 0.001, ****p ≤ 0.0001, ns=not significant.

Fig. 2: Increased HIF and ROS expression in epidermal compartment in β1 cKO skin

Quantification of temporal change in the expression epidermal HIF1a in cKO and WT skin (N=2) (A). Immunostaining of HIF1a (B) in E18.5 cKO and WT skin. Expression of GLUT1 (C), LDHa (D) and quantification (E) in chetomin treated cKO and DMSO treated cKO controls (N=2). Quantification of temporal change in the expression epidermal 8-OHdG expression in cKO and WT skin (N=2) (F). Immunostaining of 8-OHdG (G) in E16.5 cKO and WT skin (N=2). GLUT1 (H), LDHa (I) expression and quantification (J) in NAC treated cKO and PBS treated cKO controls (N=3). (K) Schematic showing glycolysis regulation by HIF1a and its regulation by ROS in cKO skin. Scale bars: 50 µm. *p ≤ 0.05, **p ≤ 0.01, ***p ≤ 0.001, ****p ≤ 0.0001, ns=not significant.

Fig. 3: Epidermis is an exporter while macrophages are importers of lactate metabolite in β1 cKO skin.

Increased membrane expression of lactate exporter MCT4 in epidermis of E17.5 and E18.5 cKO skin compared to WT (A) (N=3). Increased expression of TCA cycle enzymes CS and IDH1 (B), and quantification (C) in macrophage compartment in cKO skin compared to controls (N=2). Increase in relative levels of TCA cycle metabolites (D) and decrease in relative levels of glycolytic metabolites (E) in dermal compartment in cKO skin compared to WT (N=5). (F) Schematic showing the compartment separation of Glycolysis and TCA cycle in epidermal and macrophages compartment in cKO skin compared to WT. Increased membrane expression of lactate importer MCT1 (G, top) in macrophages in the KO skin compared to WT (N=2, n=30-40 cells each) Scale bar: 20um. Increased basement membrane (LAM332) disruption in cKO skin compared to WT (G, middle). Increased in expression on MMP9 in the cKO skin compared to WT (G, bottom). Decrease in Nuclear to Cellular ratio of expression of MCT1 in macrophages in KO and WT skin (H) (N=2, n=30-40 cells each). Quantification of basement membrane (LAM332) degradation (I) and MMP9 expression (J) in cKO skin compared to WT. (J) Schematic representing epidermis as a potential source and macrophages as potential sink for lactate metabolite in cKO skin. Acquisition of lactate by macrophage compartment correlates with increased generation of matrix remodelling enzymes that result in ECM degradation. Scale bars: 50 µm. *p ≤ 0.05, **p ≤ 0.01, ***p ≤ 0.001, ****p ≤ 0.0001, ns=not significant.

Fig. 4: Inhibition of lactate transport using small molecule inhibitor reduces pro-remodelling fate acquisition in dermal macrophages.

Decreased MMP9 expression and basement membrane (LAM332) remodelling in cKO skin treated with AZD3965, Syrosingopine compared to DMSO control (A) (N=3). Quantification of MMP9 expression in cKO skin treated with AZD3965, Syrosingopine and DMSO treated control cKO skin (B-E) (N=3). Schematic showing Syrosingopine dose schedule in imiquimod induced mice model for psoriasis (F). Decrease in epidermal plaques in Syrosingopine treated C57B6/J mice compared to 5% DMSO treated control in imiquimod induced mice model of psoriasis (G) (N=3). Reduction of epidermal thickening and number of proliferative cells (Ki67, red) in Syrosingopine treated C57B6/J mice compared to 5% DMSO treated control in imiquimod induced mice model of psoriasis (H) (N=3). Quantification of number of Ki67 positive cells in epidermal compartment (I) and epidermal thickness (J) in Syrosingopine and control treated mice treated with topical imiquimod (N=3). Quantification of MMP9 expression in dermal compartment in Syrosingopine and control treated mice treated with topical imiquimod (K) (N=3). Reduction in expression of MMP9 (L, top), number of macrophages (F4/80) (L, middle), and number of monocytes (CD11B) (L, bottom) in Syrosingopine treated psoriatic mice compared to controls (N=3). (M) Model showing that under sterile inflammatory conditions in skin, early augmentation of ROS-HIF1a axis leads to enhanced glycolysis and lactate generation in the epidermal compartment. The macrophage compartment in the dermis acts as sink of lactate released by the epidermal compartment, which is then utilized as substrate for driving TCA cycle which, in turn, is necessary for pro-remodelling fate switch macrophages. Consistently, inhibition of lactate transporters, epidermal-macrophage intrinsic metabolism and upstream glycolysis regulators in the epidermal compartment using small molecule inhibitors lead to inhibition of pro-remodelling fate and hence inflammation in the KO skin. Scale bars: 50 µm. *p ≤ 0.05, **p ≤ 0.01, ***p ≤ 0.001, ****p ≤ 0.0001, ns=not significant.

Fig. S1, related to Fig. 1: NGS data summary and validation of glycolysis and TCA cycle gene expression in epidermal compartment in β1 cKO skin.

Metabolic pathways upregulated in GSEA analysis of epidermal compartment in E18.5 cKO skin compared to WT (A). Relative transcript expression (from NGS) of glucose transporters and glycolytic genes in the epidermis of cKO and WT skin (B) (N=2). qPCR validation of glucose transporter GLUT1 and glycolytic genes in the epidermis of cKO compared to WT (C) (N=4). Quantification of temporal change in the expression of GLUT1 in cKO and WT (N=2) (D). Relative transcript expression (from NGS) of TCA cycle enzyme genes in the epidermis of cKO compared to WT (E) (N=2). qPCR validation of TCA cycle enzyme genes in the epidermal compartment compared to WT (F) (N=3). Scale bars: 50 µm. *p ≤ 0.05, **p ≤ 0.01, ***p ≤ 0.001, ****p ≤ 0.0001, ns=not significant.

Fig. S2, related to Fig. 2: Increased HIF1a target expression in epidermal compartment of the β1 cKO skin.

GSEA analysis of cKO epidermis showing upregulated HIF1a signalling and response to hypoxia at E18.5 compared to WT (A). qPCR validation of increase in transcription of downstream targets of HIF1a in the epidermis normalised to WT (B) (N=3). Epidermis of cKO skin showing increased expression of KRT14 (C, top), COX2 (C, bottom), quantified in (E), compared to WT (N=3). Epidermis of cKO skin showing decrease in expression of HIF1a targets KRT14 (D, top), COX2 (D, bottom), quantified in (G), after treatment with chetomin compared to controls (N=3). Schematic showing dose schedule of various drugs used in the study in pregnant dams carrying cKO and WT embryos (G). Scale bars: 50 µm. *p ≤ 0.05, **p ≤ 0.01, ***p ≤ 0.001, ****p ≤ 0.0001, ns=not significant.

Fig. S3, related to Fig. 2: Increased ROS response in epidermal compartment of the β1 cKO skin.

GSEA analysis of cKO epidermis showing upregulated ROS response at E18.5 compared to WT (A). cKO skin showing no significant change in the expression of 8-OHdG at E18.5 compared to WT (N=2) (B). qPCR validation of increase in transcription of both ROS source and scavenger genes, in E18.5 cKO epidermis compared to WT (C) (N=3). Epidermis of cKO skin showing decrease in expression of 8OHdG (D, top), HIF1a (D, bottom), quantified in (F), upon NAC treatment compared to controls (N=2). Epidermis of cKO skin showing decrease in the expression of KRT14 (E, top), COX2 (E, bottom), quantified in (G), upon NAC treatment compared to controls (N=3). Scale bars: 50 µm. *p ≤ 0.05, **p ≤ 0.01, ***p ≤ 0.001, ****p ≤ 0.0001, ns=not significant.

Fig. S4, related to Fig. 3: Macrophages upregulate TCA cycle genes and downregulate glycolysis in the β1 cKO skin.

Metabolic pathways upregulated in GSEA analysis of macrophage compartment in E18.5 cKO skin compared to WT (A). Relative transcript expression (from NGS) of major glucose transporters, glycolytic enzyme and TCA cycle enzyme genes in the macrophages in cKO skin compared to WT (B) (N=2). qPCR validation of glycolytic genes in cKO skin macrophages compared to WT (C) (N=3). Expression of GLUT1 (red) does not colocalize with F4/80 (green) in dermal macrophages in β1 cKO skin (D). qPCR validation of TCA cycle enzyme gene expression in cKO skin macrophages compared to WT (E) (N=3). Schematic showing macrophages as potential sinks for lactate as it can be converted to pyruvate to drive the TCA cycle (F). Scale bars: 50 µm.

Fig. S5, related to Fig. 3: Macrophages upregulate TCA cycle genes and downregulate glycolysis in the β1 cKO skin.

(A) qPCR validation of TCA cycle enzyme genes in cKO skin fibroblasts compared to WT (N=3). Metabolic pathways upregulated in GSEA analysis of fibroblast in E18.5 cKO skin compared to WT (B). (C) LAM332 staining, quantified in (D), suggest no obvious difference in the basement membrane spread in the cKO skin treated with etomoxir compared to controls. Scale bars: 50 µm.

Fig. S6, related to Fig. 4: Inhibition of Epidermal ROS and HIF1a leads to reduction in macrophage pro-remodelling fate acquisition.

Decreased MMP9 expression and basement membrane (LAM332) remodelling in cKO skin treated with TCA cycle inhibitor, PA and glycolysis inhibitor, 2DG, compared with PBS treated control cKO skin (A) (N=3). Quantification of basement membrane spread (B) and MMP9 expression (C) in PA and 2DG treated cKO and control treated cKO skin (N=3). Reduction in basement membrane spread (D, left) and MMP9 expression (D, right) in cKO skin treated with UK-5099 compared to controls. Quantification of basement membrane spread (E) and MMP9 expression (F) in cKO skin treated with UK-5099 compared to controls. Reduction in basement membrane spread in basement membrane spread (G,H,I) and MMP9 expression (G,J,K) in cKO skin treated with chetomin and NAC compared to controls. Scale bars: 50 µm. *p ≤ 0.05, **p ≤ 0.01, ***p ≤ 0.001, ****p ≤ 0.0001, ns=not significant.

Fig. S7, related to Fig. 4: Lactate mediated cross-talk in imiquimod induced psoriatic skin samples.

Increased in epidermal plaques upon induction of psoriasis using topical imiquimod for 5 days (A) (N=3). Increase in epidermal thickness and number of proliferating epidermal cells (Ki67) in imiquimod treated mice compared to Vaseline treated controls (B). Quantification of epidermal area in imiquimod treated mice compared to Vaseline treated controls (N=3) (C). Increase in macrophage (F4/80) (D, top) and monocytes (CD11B) (D, bottom) in imiquimod treated mice compared to Vaseline treated controls (N=3). Increase in MMP9 expression (E), quantified in (F), in imiquimod treated mice compared to Vaseline treated control (N=3). Increase in expression of GLUT1 (G, top) and membrane localization of MCT4 (G, bottom) in Imiquimod treated mice compared to Vaseline treated controls (N=3). Increase in expression of CS (H, top), IDH1 (H, middle) and MCT1 (H, bottom) in the macrophages in imiquimod treated mice compared to Vaseline treated controls (N=3) Scale bars: 20 µm. Quantification of GLUT1 expression in epidermis of imiquimod treated mice and Vaseline treated controls (I) (N=3). Quantification of expression of CS, IDH1 and MCT1 in macrophages in imiquimod treated mice compared to Vaseline treated controls. All experiments done for this figure are analysed at Day 6 of imiquimod treatment. Scale bars: 50 µm. *p ≤ 0.05, **p ≤ 0.01, ***p ≤ 0.001, ****p ≤ 0.0001, ns=not significant.
